# A deep learning predictor of bindable protein surfaces to guide generative synthetic biology

**DOI:** 10.64898/2026.04.16.718848

**Authors:** Leonardo Almeida-Souza

## Abstract

The advent of generative machine learning models has revolutionized *de novo* design of protein binders. However, the wide adoption of this revolution is bottlenecked by computational cost. For many targets, binder design commonly requires computationally intensive sampling across structures, often wasting days of GPU time on unwanted or geometrically inviable regions. Here, IARA (Interface Analysis and Recognition Architecture) is introduced, a deep learning Graph Neural Network designed as a rapid structural filter to triage protein binder generative pipelines. IARA is trained entirely on BindCraft trajectories generated against s RFdiffusion-generated targets. Based on a slim network with only seven residue features, IARA maps the binder designability of input proteins in a matter of seconds. On validation runs using BindCraft, RFdiffusion and BoltzGen, IARA successfully identified the optimal binding interface for practically all targets. By instantly pinpointing the highest-probability binding pockets, IARA democratizes synthetic biology, drastically reducing the exploratory GPU compute required for successful *de novo* binder generation.

## Introduction

The advent of generative structural biology marks a paradigm shift in protein engineering. A range of breakthrough tools^1–8^ allows generative targets to be rapidly and elegantly designed based on a relatively simple set of inputs. Among these tools, RFdiffusion^2^, BindCraft^1^ and BoltzGen^8^ drive one of the most promising areas in generative structural biology, the design of targeted protein binders mimicking natural ligands and inhibitors or substituting the use of antibodies. These tools require long runs on powerful GPUs and when deploying them, researchers are faced with two choices: They can either pick a region of interest in their target protein – a “hot spot” – to focus the generative efforts or allow these tools to survey the full target structure to find good binders^1,2,8^. In either case, one can end up using hundreds of hours of GPU time with no success if the hot spot or the whole protein surface is largely unfavorable for binder generation.

To truly democratize synthetic biology and mitigate these hardware bottlenecks, the field requires a fast, accurate method to prioritize generative searches *before* launching heavy GPU computations. Here, IARA (Interface Analysis and Recognition Architecture) is introduced, a Graph Neural Network built to synergize with any binder generative tool. Operating in seconds, IARA acts as a pre-computation spatial filter, directly showing protein regions most likely to yield successful *de novo* binders. IARA draws its name from Brazilian folklore as “mãe das águas” (mother of waters), a figure capable of shaping the waters and a fitting homage given the fundamental role of water in driving protein folding and protein-protein interaction.

## Results

### Training a Graph Neural Network to learn bindable protein regions

To train a machine learning model capable of recognizing bindable regions, a large dataset composed of 1067 successful BindCraft runs was generated. These runs used fully synthetic monomeric targets generated using RFdiffusion (Fig. 1a). This target selection had a few key advantages over using selected structures from pdb: i) First, it provided a “clean slate” for binder generation by avoiding any potential bias towards natural interfaces. ii) Second, it allowed easy control over the size of targets (100-250 residues in length), ensuring that each BindCraft run is within the memory constraints of the hardware (up to 550 residues for a 32GB GPU), and avoiding the need to break large proteins into smaller domains thereby exposing large hydrophobic patches. iii) Finally, it bypassed the need to curate a training dataset to avoid DNA and RNA-binding proteins or proteins containing multiple posttranslational modifications (e.g. mucins), and to remove signal peptides, transmembrane domains, amphipathic helices, and other protein features which could interfere or bias BindCraft runs.

**Figure 1.**
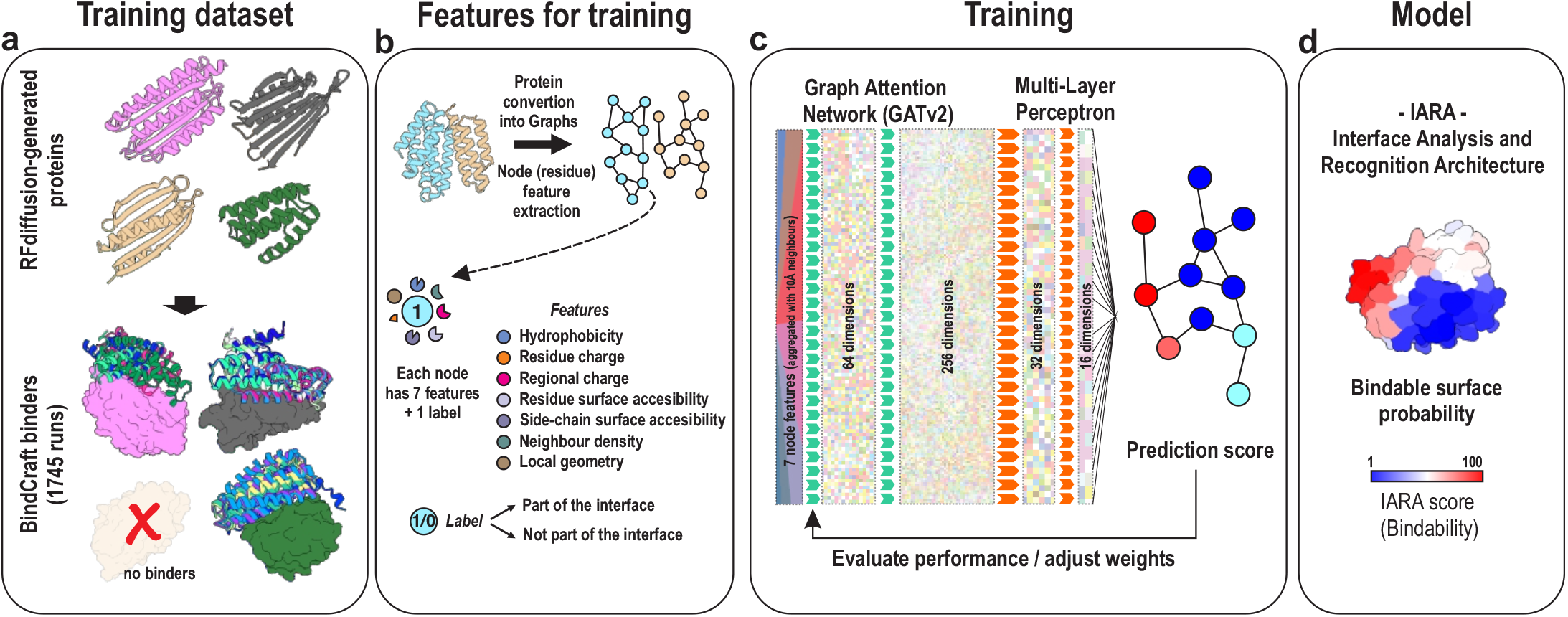
Building a graph neural network to predict bindable protein regions. **a**, Dataset generation strategy. **b**, Residue feature extraction and protein conversion to Graphs. **c**, Architecture of the neural network used for training. **d**, IARA scoring.

From these 1067 BindCraft runs, up to 5 complexes per target were selected and each protein in the complex was converted into a graph with 7 features per residue (Fig. 1b): i) hydrophobicity; ii) residue charge; iii) regional charge (within a 10Å radius), iv) residue surface-accessibility, which measures if the residue is exposed; v) side-chain surface accessibility, which measures how much of the residue side chain is surface exposed; vi) local density, which counts the number of neighboring residues within an 8Å sphere; and vii) local geometry, which measures the number of residues within a 15Å radius, acting as a proxy for curvature of the region (i.e. concave or convex). Residues on the interfaces received an “on interface” label, as the goal for the training process.

Next, the full dataset was split in training and validation sets in a 9:1 ratio. A key concern when doing this type of dataset splitting for machine learning in structural biology lies in ensuring that these sets are structurally distinct. If this structural separation is not achieved, models may produce “good” predictions on the validation dataset by simply memorizing structures from the training dataset. This phenomenon is called data leakage^9^. To assess data leakage between the training and the validation datasets, standard CAPRI metrics were employed: interface root mean squared deviation (iRMSD) and the fraction of native contacts (F_nat). All 107 targets in the validation set were compared to all complexes in the training set, totaling > 600.000 comparisons. For 106 out of 107 validation targets (>99%), the minimum average iRMSD was 2.6Å with maximum F_nat values primarily between 0.3 and 0.7 (Supplementary fig. 1). According to CAPRI criteria, these values represent ‘Acceptable’ to ‘Medium’ structural similarity, indicating a shared physical binding grammar (e.g., common secondary structure packing, which is expected for protein-protein interactions) rather than atomic-level identity. Only one target exhibited a significant similarity (iRMSD=2.2Å, F_nat = 0.84) to one of the complexes in the validation set. This target was kept in the validation dataset, acting as a control to check for model bias towards it. Thus, the absence of systemic structural overlap demonstrates that the validation and training sets represent a distinct pool of target-binders.

With the datasets defined, the training set was fed into a 3-layer Graph Attention Neural Network^10^ and trained for 150 epochs. During training, two metrics were evaluated: the loss in the validation dataset and signal to noise ratio (SNR = maximum score in a protein interface / mean surface score). The final model used in the rest of this study achieved a validation loss of 0.1952 and a signal-to-noise ratio SNR of 2.43 (Fig. 1c).

This final model was named IARA (Interface Analysis and Recognition Architecture) (Fig. 1d). The model takes as input any protein structure format (pdb, cif, pdb.gz, and ent), calculates the IARA score (“*bindability*”) for each residue (from 0 to 100), and writes the score to the b-factor column of an output pdb file. The model takes a few seconds to run, does not require a GPU and can be used as a standalone python script, a Pymol or Chimerax plugin or via a Google Colab notebook. All these usage modalities are available to download at GitHub: https://github.com/leodeals/IARA.

### IARA successfully predicts bindable surfaces for BindCraft and outperforms MaSIF in this task

To confirm that the model successfully learned the core principles of BindCraft’s generative preferences, IARA was evaluated on a validation set of 107 unseen synthetic targets and their corresponding 532 generated binders. For this analysis, three IARA score thresholds (IARA > 50, > 60 and > 70) were tested. For each of these thresholds, it was measured if an IARA patch above the threshold was in contact – within 5Å – with a binder and what percentage of the target interface had IARA scores above any given threshold. These measurements indicated if IARA could not only correctly predict a binder interface, but also how much of the binding interface it could pinpoint (predicted interface accuracy). These measurements were done at the target level (has IARA identified the region(s) in the target protein where binders were generated?) and on the binder level (has IARA identified these regions *for all* binders generated?) At the target level, IARA correctly identified binding interfaces for 100%, 96%, and 92% of the 107 targets at the > 50, > 60, and > 70 thresholds, respectively (Fig. 2a and Supplementary table 1). When evaluating at the binder level, IARA accurately predicted the interaction footprint for 99%, 97%, and 87% of the 532 generated binders across the respective thresholds (Fig. 2b, c and Supplementary table 2). Further confirming that IARA has learned surface bindability rather than memorized a few examples via data leakage, the target showing the highest similarity with the training dataset in the interface comparison analysis shown in supplementary fig. 1a (D10_1070) ranked only 91^st^ out of 107 in predicted surface accuracy (Supplementary table 1).

**Figure 2.**
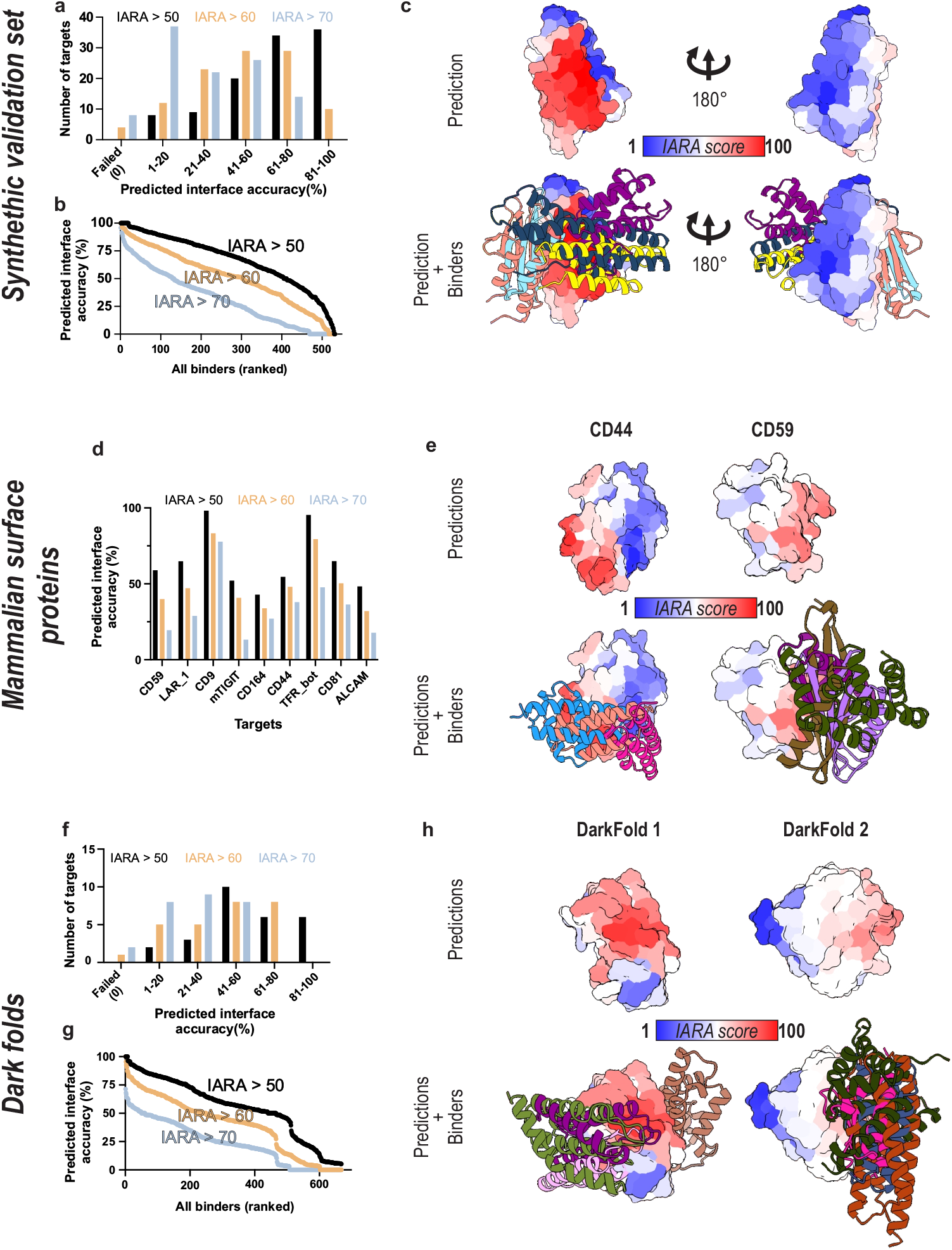
IARA predictions for BindCraft. **a**, Frequency distribution of the predicted interface accuracies for targets in the synthetic validation dataset. **b**, Predicted interface accuracies for all binders generated for the synthetic validation dataset. **c**, Example of a structure from the synthetic validation dataset (as surface) showing the IARA prediction scores from blue (low bindability) to red (high bindability). The structure of five binders are shown as colored ribbons. **d**, Frequency distribution of the predicted interface accuracies for each mammalian surface protein tested. **e**, CD44 and CD59 structure showing the IARA prediction scores. The structure of three and four binders for each protein, respectively, are shown as colored ribbons. **f**, Frequency distribution of the predicted interface accuracies for “dark fold” targets. **g**, Predicted interface accuracies for all binders generated for “dark fold” proteins. **h**, Structures of dark fold 1(D103^12^) and dark fold 2 (D29^12^) showing the IARA prediction scores. The structure of five and four binders for each protein, respectively, are shown as colored ribbons. Analyses were performed at three IARA thresholds > 50 (black symbols and bars), > 60 (orange symbols and bars) and > 70 (cyan symbols and bars).

To contextualize IARA’s performance to define bindable surfaces, its predictions were benchmarked against MaSIF^11^, a pioneering geometric deep learning framework widely used for identifying generalized protein interaction surfaces. While MaSIF excels at identifying broad, naturally occurring interaction patches for protein-protein and protein-ligand interactions, it requires computationally intensive steps. Therefore, it was tested if IARA could serve as an alternative, faster, and streamlined tool with the precision required for rapid evaluation of protein surfaces. Interestingly, IARA demonstrated significantly higher sensitivity for predicting binding hot spots. While IARA recovered anchoring sites for 100% of the validation targets at the > 50 threshold, MaSIF recovered hotspots for 46.7% of targets under identical conditions, capturing an average of just 4.6% of the interface residues (Supplementary fig. 2a, Supplementary tables 3 and 4). These results suggest that while general tools like MaSIF remain valuable for broad interface characterization, the specialized graph-based architecture of IARA provides the targeted precision necessary for successful *de novo* binder generation.

To further confirm the usability of IARA, the same analysis was performed using BindCraft runs for 9 mammalian cell surface proteins (ALCAM, CD59, CD9, CD164, CD44, CD81, TIGIT, LAR1, and TFRC). For these proteins, transmembrane domains and secretion signals, when present, were removed prior to computational inference. For the TFRC (transferrin receptor), the extracellular region was split into top and bottom (bot) domains. The top domain did not result in any binders, so it was excluded from this analysis. All proteins were run on BindCraft without the definition of specific “hot spots”, to ensure a full exploration of the protein surfaces.

Despite being trained exclusively on synthetic targets, IARA achieved perfect precision with these “real-world” examples at the target level, with 100% successful binding interfaces identified across all three thresholds (Fig. 2d, e and Supplementary table 5). At the binder level, success rates were 100%, 100%, and 95% (out of 457 binders, Supplementary Fig. 2d and Supplementary table 6). Thus, IARA can confidently predict bindable surfaces preferentially used by BindCraft and shows better predictability for this task than MaSIF.

### IARA successfully predicts bindable surfaces in unnatural protein folds

One of the most interesting uses of generative synthetic biology lies in the possibility to create novel proteins with novel properties and unusual folds^12^. To test if IARA’s understanding of bindable protein surfaces can also be used for non-standard proteins, we tested its performance against a dataset of “dark folds”. These proteins were designed with the explicit goal to target unexplored regions in the protein fold space^12^.

To test this possibility, BindCraft binders were generated for 27 available dark folds^12^ and IARA’s performance was evaluated against the results. Confirming that IARA can predict bindable surfaces in any protein fold, at the target level, IARA pinpointed the correct bindable surface for 27 (100%), 26 (96%), and 25 (93%) targets at the > 50, > 60, and > 70 thresholds, respectively (Fig. 2f and Supplementary table 7). At the binder level, it predicted the interaction footprint for 100%, 90%, and 66% of the 668 generated binders(Fig. 2g, h, and Supplementary table 8). Taken together, IARA shows remarkable predictive capacity for bindable protein surface for BindCraft runs using synthetic, natural and unusual protein folds.

### IARA successfully predicts bindable surfaces for RFdiffusion and BoltzGen

Given the remarkable IARA’s performance finding bindable surfaces for BindCraft, it was hypothesized that the principles learned by the model could also be applicable for RFdiffusion and BoltzGen binder design (Fig. 3a). For that, binders for the synthetic validation set and the mammalian surface proteins were generated using these pipelines and their results compared to IARA predictions.

**Figure 3.**
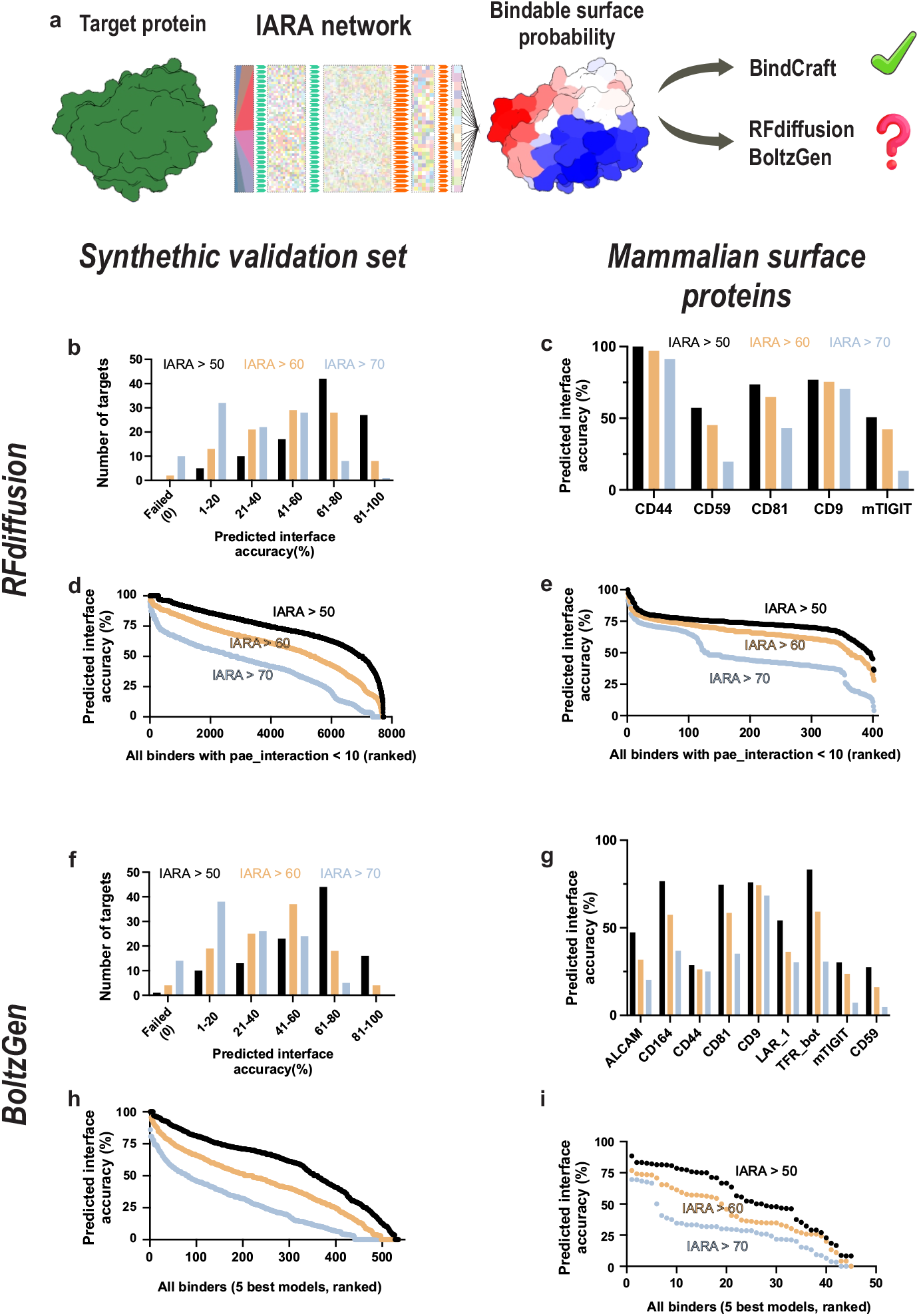
IARA predictions for RFdiffusion and BoltzGen-generated binders. **a**, Can a model trained on BindCraft data also predict bindable surfaces for RFdiffusion and BoltzGen? **b, f** Frequency distribution of the predicted interface accuracies for targets in the synthetic validation dataset ran on RFDiffusion (b) and BoltzGen (f). **c, g** Frequency distribution of the predicted interface accuracies for a set of mammalian surface proteins ran on RFdiffusion (c) and BoltzGen (g). **d, h** Predicted interface accuracies for all RFDiffusion (d) and BoltzGen (h)-generated binders for the synthetic validation dataset. **e, i** Predicted interface accuracies for all RFDiffusion (e) and BoltzGen (i)-generated binders for a set of mammalian surface proteins. Analyses were performed at three IARA thresholds > 50 (black symbols and bars), > 60 (orange symbols and bars) and > 70 (cyan symbols and bars).

When compared to BindCraft results, IARA achieved a similar level of precision with binders generated with RFdiffusion and BoltzGen (Fig. 3b-i). At the target level, IARA predicted the binder surfaces in the synthetic validation dataset for 100%, 98% and 91% of targets for RFdiffusion runs and 99.1%, 96.3% and 87.9% of targets for Boltzgen runs (Fig. 3b,f and Supplementary tables 9 and 13). For the mammalian surface proteins, IARA successfully predicted the bindable surfaces for all targets which generated successful binders for both RFdiffusion and BoltzGen across all three thresholds (Fig. 3c, g and Supplementary tables 10 and 14).

At the binder level, IARA predicted the interaction surface for all RFdiffusion generated binders for the mammalian surface proteins dataset (of 1206 binders) and 99.6%, 99.3%, and 95.1% for the synthetic proteins dataset (7,733 binders) at the > 50, > 60, and > 70 thresholds, respectively (Fig 3d, e and Supplementary tables 10 and 12). For BoltzGen, IARA predicted 100%, 97.8% and 93.3% of all binders (out of 45 binders) for the for the mammalian surface proteins dataset and 98.5%, 92.9% and 82.2% for the synthetic proteins dataset (535 binders) (Fig 3h, i and Supplementary tables 15 and 16). Taken together, IARA is a tool capable of predicting bindable surfaces for all commonly used binder generation tools.

## Conclusion

The discrepancy between what generative machine learning tools are capable of designing, and what researchers can afford to compute and experimentally validate, represents a significant friction point in modern synthetic biology. IARA bridges the computational side of this gap. By utilizing a light Graph Neural Network trained on only 4 fundamental proteins characteristics spread across 7 parameters, it provides an ultra-fast structural triage tool that tells researchers exactly where to aim their computational resources for all commonly used generative binding tools available: BindCraft, RFdiffusion and BoltzGen^1,2,8^.

The apparent imbalance between IARA’s performance with the size of its training dataset and the depth of its GNN stresses a fundamental reality in the machine learning field^13,14^. High quality datasets can provide better results than large, noisy datasets. Efforts to curate or generate high quality datasets can be instrumental to tackle the biggest challenges in computational biology.

It is important to stress that IARA cannot distinguish between binders which will work experimentally and the ones which will not. This would require detailed binding data for thousands of targets and binders, which is currently not available at the scale required. However, given the relatively high success rates reported for BindCraft (20-80%), IARA represents a crucial step in democratizing synthetic biology in times of increasing costs for computing power.

## Materials and Methods

### Dataset generation, feature extraction and training

The targets for the training dataset were generated using RFdiffusion for backbone generation ranging between 100-250 residues, followed by sequence optimization via ProteinMPNN, and final structural prediction via AlphaFold2. Approximately 10,000 total targets were generated and only targets exhibiting an AlphaFold2 predicted Local Distance Difference Test (pLDDT) score > 0.92, totaling 1745, were used for downstream processing.

These targets were then run through the default 4-stage BindCraft pipeline to generate successful binding trajectories. The training dataset aggregated outputs from two distinct run modes: 774 successful 1-day runs (out of 1,346 attempts) and 293 successful 3-day runs (out of 399 attempts). For these runs, the number of final successful designs were capped to 5 for up to 1-day runs and 100 for up to 3-day runs. All runs were performed on Nvidia Volta V100 graphics cards on the Puhti supercomputer at the Finnish IT Center for Science (CSC). Together, these runs represented over 57,800 hours of GPU usage.

From these successful BindCraft runs, up to 5 geometrically successful target-binder complexes per individual target were selected (to avoid overrepresentation of 3-day runs). This selection yielded 3,785 unique protein-protein complexes and each side of the interaction surface was used to compute local features, totaling a dataset of 7,570 distinct structural interaction footprints. The full structurally validated complexes were strictly split 9:1ratio at the *target* level into training and validation sets.

For each interaction footprint, 7 features were extracted:

1. Hydrophobicity: Kyte-Doolittle scale was used.
2. Residue charge: Assigned directly from standard amino acids properties. +1 for Arginine and Lysine, −1 for Aspartate and Glutamate and 0 for the remaining 16 standard amino acids.
3. Charge patch. Evaluates the cumulative electrostatic environment by assessing neighboring carbon α atoms within a 10Å sphere and dividing by 0.12 to scale this feature with others. This scaling kept the variance of this feature around zero, a common trick to improve model convergence^15^.
4. Residue surface exposure. Estimates residue surface localization based on the half-sphere exposure metric^16^. This calculates the number of carbon α neighbors within 8Å (dens8) and applies the formula 1 - (dens8 / 100).
5. Residue surface accessibility (i.e. relative surface exposure). Estimates side-chain exposure by dividing the unscaled density-based surface exposure by the reference maximum surface area of that specific residue side chain.
6. Local density. Quantifies local structural crowding by counting the number of carbon α neighbors within 8Å and dividing by 10.
7. Local geometry. Assesses broader, lower-resolution protein geometry by counting the number of carbon α neighbors within 15Å and dividing by 10. In addition to these features, each protein residue was assigned a label, signaling its presence on or off a protein interface.

The network was trained for exactly 150 epochs using the Adam Optimizer (initial Learning Rate = 0.001, Weight Decay = 1e-4) equipped with a “ReduceLROnPlateau” scheduler.

The training algorithm utilized a 3-layer Graph Attention Network (GATv2) architecture to synthesize local physicochemical information across the target protein surface. In the initial layer, each amino acid aggregates features from neighbors within a 10Å radius, projecting the 7 initial physical features. This layer is then passed to the next two layers with high-dimensional hidden representations (progressing from 64 to 256 dimensions). This projection allows the capture of complex, non-linear spatial relationships. In the final graph layer, this rich neighborhood context is compressed into a refined 32-dimensional node embedding. These highly contextualized embeddings were subsequently processed by a Multi-Layer Perceptron (MLP) head, which reduces the 32 dimensions down to 16 dimensions and finally to a single raw interaction prediction. Finally, this raw score is mapped to a normalized 0-to-100 probability (the IARA score) via a Sigmoid activation function.

### Comparison of validation and training sets

To rigorously evaluate the structural novelty of the validation dataset and quantify potential data leakage from the training set, a sequence-independent, 3D structural alignment pipeline inspired by the Critical Assessment of Predicted Interactions (CAPRI) criteria was implemented.

Interface Definition and Coordinate Extraction: For every protein complex, the binding interface was isolated using the relative spatial positioning of the carbon-α atoms. An interface was defined by the subset of carbon-α atoms from the Target (chain A) and Binder (chain B) that resided within a 10Å radius of one another. The neighborhood search was implemented using scipy.spatial.cKDTree ^17^, while structural parsing was handled using the ProDy library^18^.

Spatial Alignment: Rigid-body alignments were performed using the Iterative Closest Point (ICP) algorithm^19^. ICP aligns two unlabeled point clouds by iteratively minimizing the point-to-point distance between them. At each iteration, nearest-neighbor pairings were re-established utilizing k-d trees. This iterative process was executed using numpy array operations with a maximum limit of 20 iterations and a strict convergence tolerance (stopping when the change in average distance between iterations was < 0.001Å).

Evaluating CAPRI Metrics: Once the optimal rotation and translation operators were established by aligning the isolated interface subsets of the fixed targets, the entire interface complex was transformed into the validation coordinate space. Two continuous variants of conventional CAPRI statistics to assess structural homology were calculated:

1. Interface Root Mean Square Deviation (iRMSD): A sequence-independent spatial iRMSD was calculated over the complete interfacial point cloud (target plus binder) using the formula:

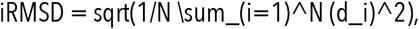 Where N is the total number of interface atoms, and d_i is the nearest Euclidean distance between the i-th aligned training atom and its closest validation atom.
2. Spatial Fraction of Native Contacts (f_nat): To quantify the specific degree to which a binder’s spatial pose was replicated by a training set analog, a purely spatial adaptation of F_nat was calculated. The mobile subset of the interface (e.g., the binder) was projected into the aligned pose and a contact was considered successfully “reproduced” if a training binder carbon-α atom fell within an overlap distance threshold of 2Å of any validation binder carbon-α atom using the formula:

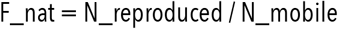

where N_reproduced represents the number of binder atoms falling within the alignment threshold, and N_mobile is the total number of binder interface atoms in the validation complex.

Every unique validation target was exhaustively queried against the full training set using direct (Target vs Target) and flipped orientation (Target vs Binder). Pairs yielding the maximum spatial f_nat and minimum iRMSD metrics were recorded as the nearest conservative measure of cross-set data leakage.

### Evaluation datasets & Bindcraft, RFdiffuion and BoltzGen runs

The synthetic validation dataset represents a 10% holdout from the initially generated dataset pool. It is composed of 107 RFdiffusion generated targets which were run on BindCraft (for either 1- or 3-days). The same targets used for BindCraft were also used for binder generation using RFdiffusion and Boltzgen.

*RFdiffusion* runs were setup using the RFdiffusion-proteinMPNN-AF2 binder generation pipelines using the standard settings (as described in the RFdiffusion GitHub page) without defining hot spots. 500 designs per target were generated and filtered for RoseTTAfold^20^ pae_interaction (predicted aligned error for the interaction surface) score below 10. This is the suggested threshold to select designs worthwhile to be used in experimental validation.

*BoltzGen* runs were setup using the “protein-anything” protocol with standard settings without defining hot spots. In this case 4000 designs were generated and the best 5 designs (scored by the internal scoring matrix of BoltzGen) were selected for comparison with IARA scores.

The mammalian surface proteins dataset is composed of 8 human proteins (ALCAM, CD59, CD9, CD164, CD44, CD81, TFRC and LAR1) and one mouse protein (Tigit). The table below shows the details for each target and their respective BindCraft and RFdiffusion runs. As described above, all BoltzGen runs generated 4000 designs and the 5 best ones were picked.

| Target name | Structure ID (residues) | BindCraft<br># accepted | BindCraft GPU<br>time | RFdiffusion<br># total designs/ #pae_int <10 |
| --- | --- | --- | --- | --- |
| **ALCAM** | 5A2F (full, no NAGs) | 24 | 288h | 2000/0 |
| **CD59** | 2UWR (full) | 184 | 360h | 500/35 |
| **CD9** | 6K4J (full) | 2 | 360h | 500/111 |
| **CD164** | AF-Q04900-F1-model_v4 (57 to 111) | 17 | 288h | 2000/0 |
| **CD44** | 1UUH (full, monomer only) | 5 | 288h | 500/1 |
| **CD81** | 1G8Q | 93 | 360h | 500/242 |
| **LAR1** | 2YD8 (30-227) | 61 | 288h | 2000/0 |
| **Tigit** | AF-P86176-F1-model_v6 (26-131) | 115 | 288h | 500/13 |
| **TFR (bot)** | 1CX8 (198-379) | 2 | 360h | 2000/0 |

The dark folds dataset^12^ was downloaded from Zenodo. All 27 pdbs from this dataset were run on BindCraft for 3 days walltime (or 100 designs) using the standard settings without defining hot spots.

### MaSIF runs

To generate binding site predictions using the MaSIF-site architecture, all the 107 targets in the validation dataset were run through the three-phase MaSIF pipeline. In Phase 1 (Surface Triangulation), solvent-excluded surfaces were computed using MSMS (Michel Sanner’s Molecular Surface) with a standard 1.5 Å water probe radius to generate a dense triangulated geometric mesh of the protein exterior. In Phase 2 (Feature Extraction), the mesh resolution was down-sampled to 1.0 Å spatial clustering to optimize memory allocation, and biochemical and topological parameters were mapped to every vertex. In Phase 3 (Neural Inference), the annotated surface was segmented into overlapping circular geodesic patches which were subsequently convolved through a 3-layer Geometric Deep Neural Network to predict a continuous probability of protein-protein interaction propensity for each individual local patch. The results from the MaSIF-site run were compared against IARA using the metrics shown below. In addition to MaSIF score >50 threshold, the analysis was also done with lower and higher thresholds (supplementary tables 3 and 4).

### Calculation of validation metrics

To quantitatively assess IARA’s predictive accuracy against generated binders, two primary validation metrics were established: Target Hit Rate and Predicted interface accuracy. For each generated complex, the ground-truth physical interface was defined as any target protein carbon α atom residing within 5Å of any binder atom.

Target Hit Rate (Binary Success): A generated binder was classified as a ‘Hit’ if at least one residue within its ground-truth interface was correctly predicted as a hotspot by a defined IARA threshold (> 50, > 60 and > 70). The target-wise hit rate is the percentage of successful designs meeting this criterion.

Predicted interface accuracy: To evaluate the extent of IARA’s spatial guidance, the fraction of the ground-truth interface that overlapped with IARA predictions was calculated. This was defined as the percentage of interfacial target residues that yielded an IARA predicted probability above each threshold.

### Software Availability

To maximize accessibility for all researchers, IARA is deployed across multiple environments. The tool is available as a standalone command-line script, as native interactive graphical plugins for both PyMOL and UCSF ChimeraX, and as a zero-install Google Colab notebook at https://github.com/leodeals/IARA.

## Acknowledgements

I would like to thank the Finnish IT Center for Science (CSC) for providing and maintaining the computational resources used in this project. I am also thankful for the CSC staff for help with installing and debugging of packages.

A special thank you to Markku Hakala, Illona Rissanen & Ville Paavilainen for the critical and kind reading of the manuscript. I also would like to thank all the members of my lab for discussions during this project.

LA-S is supported by HiLIFE & the Research Council of Finland (Research Fellow).

## Supplementary figures

**Figure S1.**
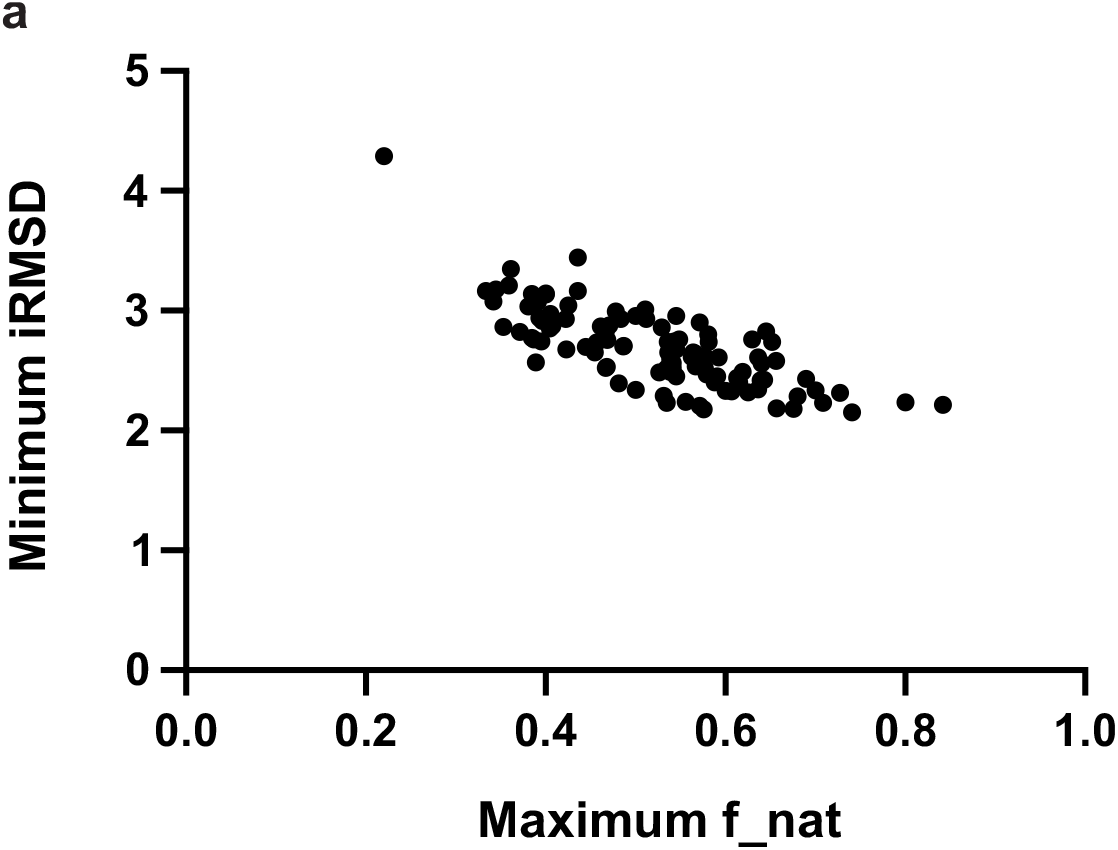
The validation and training datasets are structurally distinct. **a**, Graph displaying the minimun iRMSD and maximum f_nat scores for each one the 107 targets in the validation dataset. Note that these are the poorest scores for each target, among the >600.000 comparisons made.

**Figure S2.**
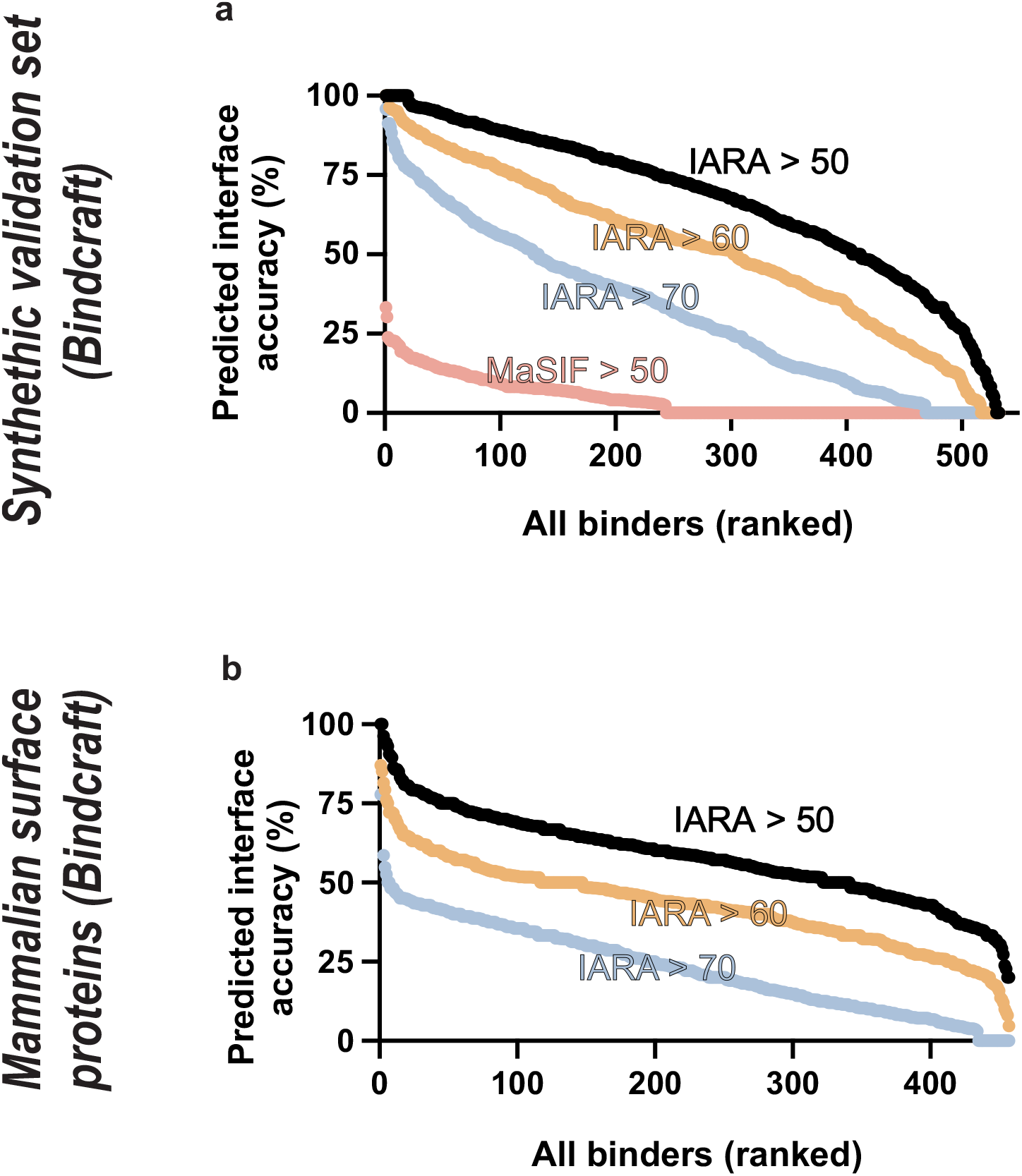
Extended info on IARA predictions for BindCraft. **a**, Comparison of the IARA and MaSIF predicted interface accuracies for all for targets in the synthetic validation dataset. **b**, Predicted interface accuracies for all for targets in the mammalian surface proteins set. Analyses were performed at three IARA thresholds > 50 (black symbols and bars), > 60 (orange symbols and bars) and > 70 (cyan symbols and bars). For MaSIF analysis, a >30, >40, >50, >60 and >70 was performed (Supplementary tables 3 and 4).

